# Methodological Evaluation and Data Resource for Andes Virus Sequencing Preparedness

**DOI:** 10.64898/2026.05.15.725146

**Authors:** Rebecca Doherty, Kuiama Lewandowski, Abigail Fenwick, Isobel Everall, Daniel Morley, Hassan Hartman, Siobhan Staplehurst, Christopher Kent, Nicholas J. Loman, Joshua Quick, Steven T. Pullan

## Abstract

As part of preparedness activities supporting pathogens classified under the UK High Consequence Infectious Diseases (HCID) framework, we previously evaluated both a whole-genome tiling amplicon sequencing scheme and a pan-viral hybridisation capture approach (TWIST-CVRP) for sequencing Andes virus (ANDV).

In light of the recent outbreak, we make available viral sequencing datasets generated using a historical ANDV isolate (Chile, 1997). In addition, we provide an evaluation of tiling amplicon scheme performance and present recommended primer updates informed by *in silico* comparison with the recently released outbreak genome. These datasets are intended to support benchmarking, validation, and optimisation of bioinformatic pipelines across the community.

## Background

Andes virus (ANDV), a New World hantavirus, is a high-consequence zoonotic pathogen responsible for hantavirus pulmonary syndrome and notable for its capacity for limited person-to-person transmission. Despite its clinical importance, sequencing approaches for ANDV are less mature than for many other RNA viruses.

Preparedness efforts require both robust wet-lab protocols and validated bioinformatic pipelines. Publicly available, well-characterised sequencing datasets are critical for enabling cross-laboratory benchmarking and pipeline validation, particularly for pathogens with limited routine sequencing.

Here, we provide a curated sequencing dataset and methodological evaluation derived from a historical isolate, alongside updated guidance for amplicon-based sequencing of the current outbreak viral genome.

We release FASTQ files (viral-mapped reads only) from:

- Illumina data derived from tiling amplicon sequencing
- ONT TWIST-CVRP sequencing data

These datasets are intended for:

- Bioinformatic pipeline validation
- Method comparison
- Training and benchmarking exercises

Accession details and repository links:

https://quick-research-andv-data.s3.climb.ac.uk/andes_amplicon_scheme_r1.fastq.gz

https://quick-research-andv-data.s3.climb.ac.uk/andes_amplicon_scheme_r2.fastq.gz

https://quick-research-andv-data.s3.climb.ac.uk/andes_twist.fastq.gz

## Methods

### Source Material

RNA was extracted from a 1997 Chilean Andes virus isolate (9717869/AF291704) (1) obtained from the European Virus Archive GLOBAL (EVA-G, https://www.european-virus-archive.com/), originally submitted by the Bernard-Nocht Institute of Tropical Medicine (Hamburg, Germany).

### Tiling amplicon sequencing

A 1 kb tiling amplicon scheme (ukhsa-andes/1000/v1.0.0) was designed using VarVAmp v1.2.2(2). MAFFT (3) Alignments of publicly available genomes for each segment (comprised of 48 L, 67 M and 83 S segments, supplementary file S1.) were used as input with threshold setting of 0.8, maximum of 3 ambiguous bases, optimum length of 1 kb, maximum length of 1.5 kb and minimum overlap of 100bp.

The S segment alignment included two sequences approximately 587 bp longer than others. To ensure amplification of genomes that potentially did not include this additional length, an extra primer pair was manually designed and added to the scheme. This resulted in a total of 14 primer pairs distributed across three pools.

cDNA synthesis and multiplex PCR amplification were performed following the ARTIC SARS-CoV-2 sequencing protocol v4 (4). PCR products were pooled and sequenced on the Illumina NextSeq 2000 platform, using an Illumina DNA prep library. Reads were mapped to the Andes virus RefSeq genome for each segment (NC_003468.15, NC_003467.2, NC_003466.1) using BWA-MEM (5), and coverage statistics generated using SAMtools (6).

Full details of the scheme are available at primalscheme labs (labs.primalscheme.com/detail/ukhsa-andes/1000/v1.0.0)

### In silico assessment of suitability of the amplicon scheme to the latest outbreak strain

The release of the ANDV/Switzerland/Hu-3337/2026 genome (7) (Genbank: PZ385161, PZ385162, PZ385163) enabled an *in-silico* assessment of the performance of the existing scheme. PrimalScheme3.3.0 was used to find mismatches between the primers and the ANDV/Switzerland/Hu-3337/2026 genome sequence. A development model was used to predict the effect of single primer mismatches on amplification efficiency for Q5 DNA polymerase (NEB).

### Twist Comprehensive Viral Research Panel and ONT sequencing

cDNA synthesis, library preparation, and target enrichment by hybridisation were performed using the Twist Library Preparation EF Kit 2.0 for ssRNA virus detection, with PCR conditions modified to promote longer fragment generation. Enriched libraries were subsequently prepared for Oxford Nanopore sequencing using the Ligation Sequencing Kit V14 (SQK-LSK114) protocol. Sequencing was carried out on a GridION platform using R10 flow cells according to the manufacturer’s standard protocol. Reads were mapped to the Andes virus reference genome (AF291704) using minimap2 (8).

## Results

The tiling amplicon scheme ukhsa-andes/1000/v1.0.0 achieved near-complete genome recovery across all three ANDV segments. A single amplicon within the L segment showed reduced amplification efficiency, resulting in a localised coverage drop (Figure 1).

**Figure 1.**
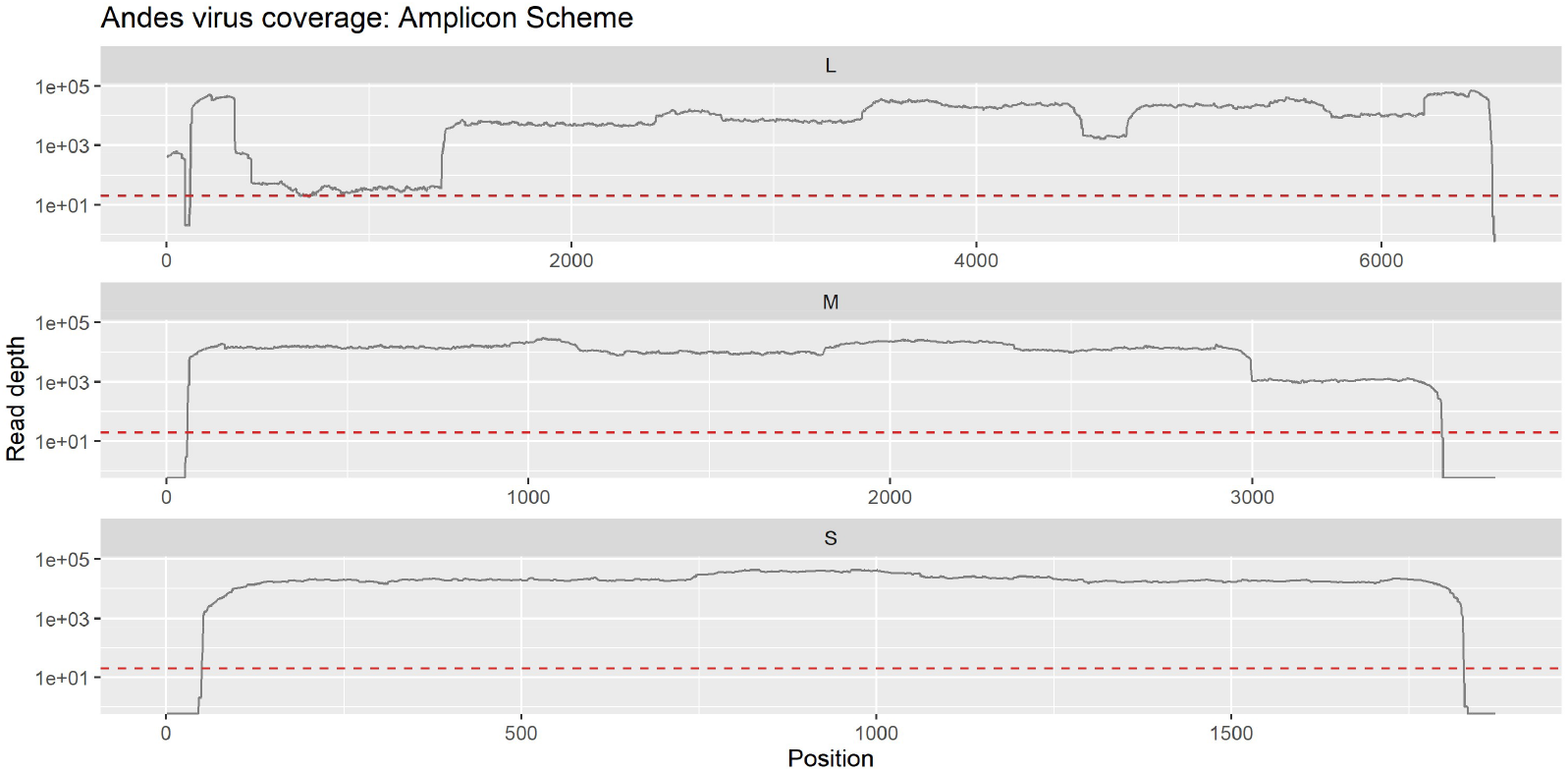
Genome segment coverage obtained using the 1 kb tiling amplicon sequencing scheme. Read depth across each viral genome segment is plotted following mapping to the reference genome. The red horizontal line indicates a depth of 20× coverage.

TWIST-CVRP bait-capture sequencing enabled uniform recovery of full-length consensus sequences across all three genome segments, with coverage exceeding 20× throughout (Figure 2).

**Figure 2.**
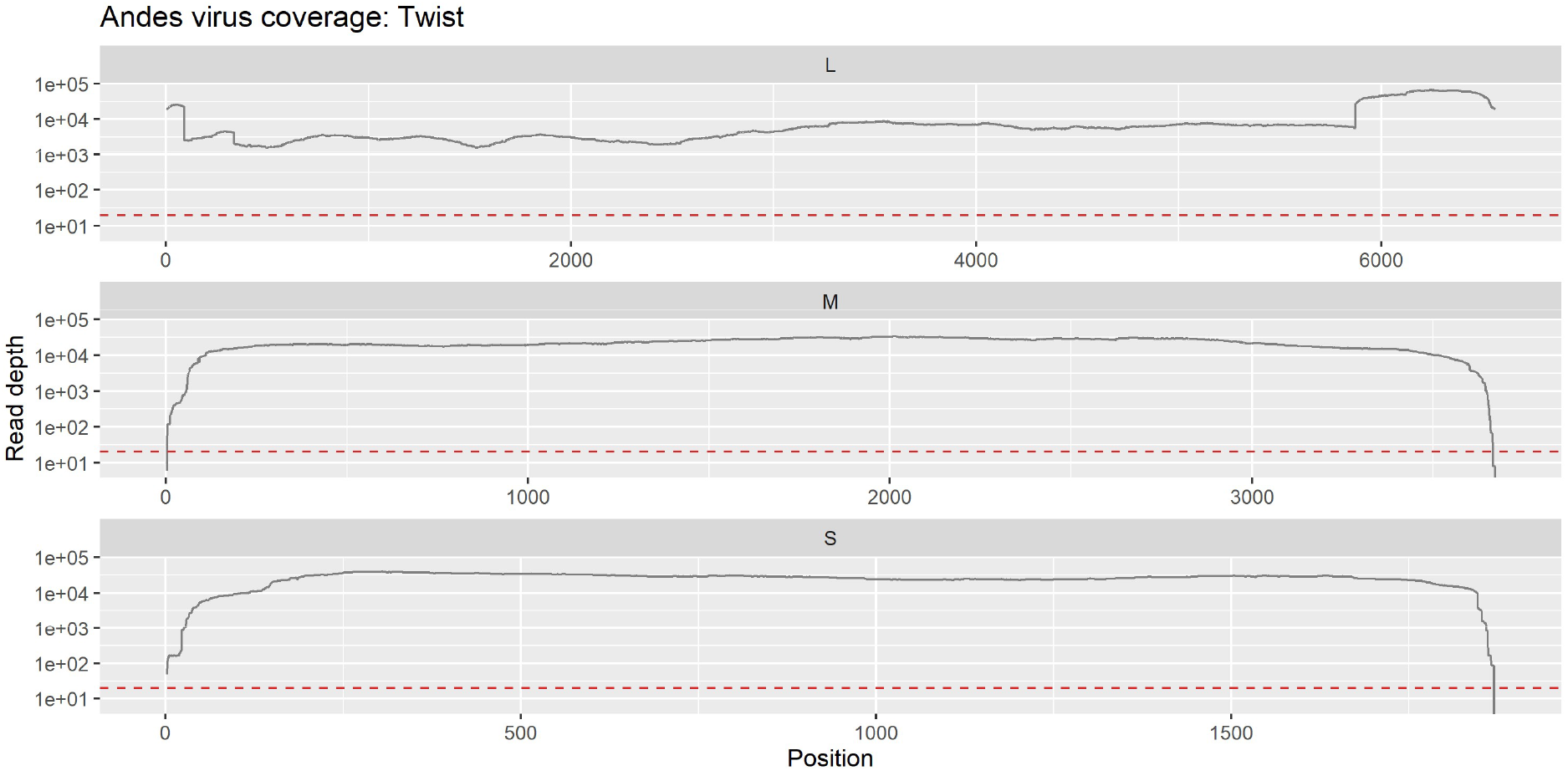
Genome coverage generated from TWIST-CVRP bait capture sequencing. Mapped read depth achieved across each segment shown, red line indicates a depth of 20x coverage.

### Updated Primer Scheme Assessment

4 replacement primers were generated to replace those with significant delta cycle threshold (dCt < -2) predicted for mismatches or those with multiple mismatches (Figure 3). Two redundant primers (Andes_S_trim_1000_0_LEFT_V2, Andes_S_trim_1000_0_RIGHT) were removed to produce a modified scheme ukhsa-andes/1000/v1.1.0 (detailed at labs.primalscheme.com/detail/ukhsa-andes/1000/v1.1.0).

**Figure 3.**
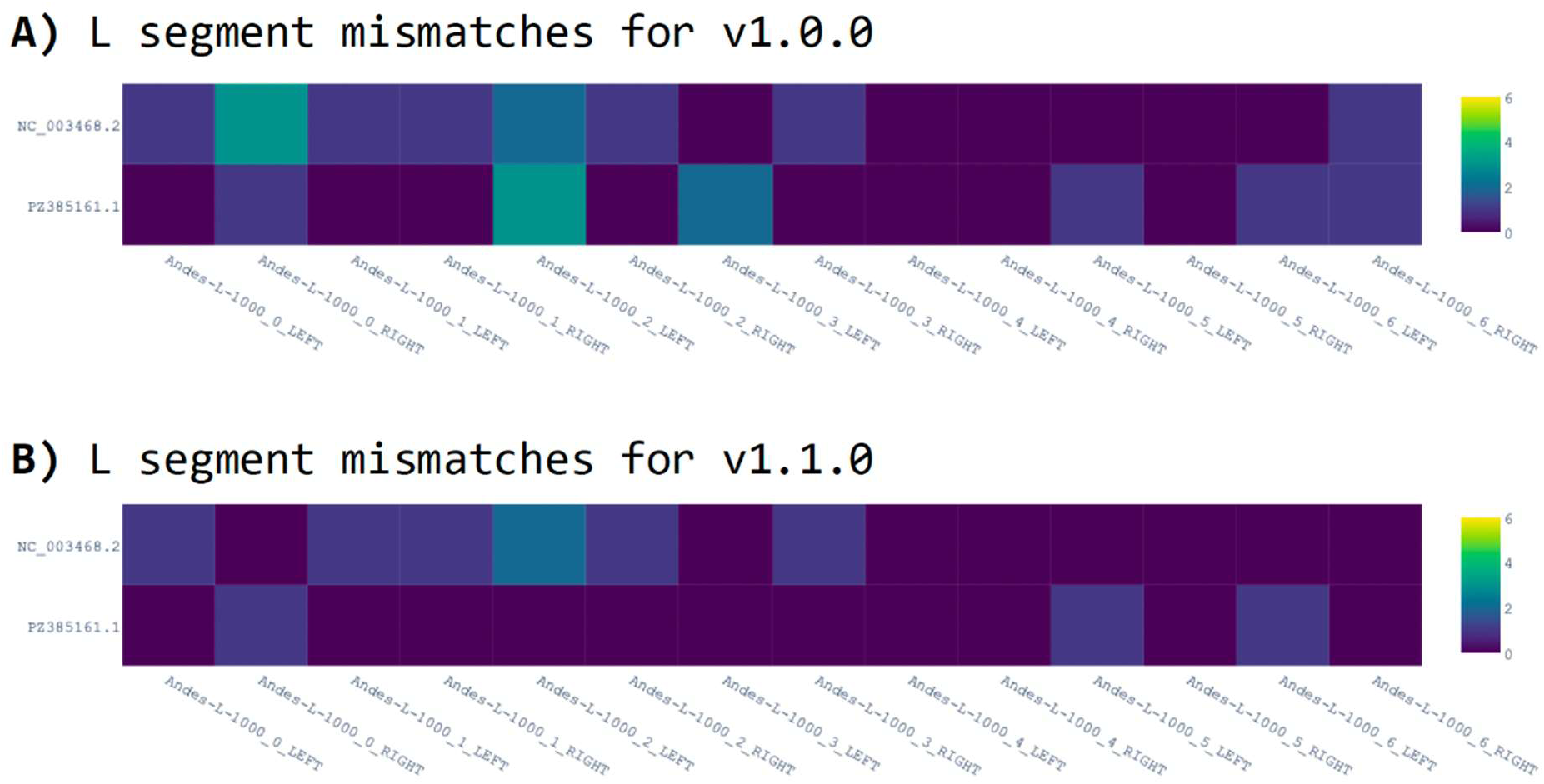
L segment mismatch profile for original and modified primer schemes.

## Conclusion

We provide a publicly available sequencing dataset and updated methodological guidance to support the global response to Andes virus.

By enabling pipeline validation and improving primer scheme design, this work contributes to strengthened genomic preparedness and more consistent analytical practices across laboratories.

## Supporting information

S1 - List of references sequences used in MAFFT alignment

## Supplementary Files

S1 - List of references sequences used in MAFFT alignment

## Acknowledgements

This study has been funded in part through the National Institute for Health and Care Research (NIHR) Health Protection Research Unit in Public Health Genomics. The views expressed are those of the author(s) and not necessarily those of UKHSA, the NIHR or the Department of Health and Social Care.”

